# ARPC5 deficiency leads to severe early onset systemic inflammation and early mortality

**DOI:** 10.1101/2023.01.19.524688

**Authors:** Elena Sindram, Andrés Caballero-Oteyza, Naoko Kogata, Shaina Huang, Zahra Alizadeh, Laura Gamez-Diaz, Mohammad Reza Fazlollhi, Xiao Peng, Bodo Grimbacher, Michael Way, Michele Proietti

**Affiliations:** Institute for Immunodeficiency, Center for Chronic Immunodeficiency, Medical Center, Faculty of Medicine, Albert-Ludwigs-University, Freiburg, Germany; Spemann Graduate School of Biology and Medicine (SGBM), Albert-Ludwigs-University of Freiburg, Germany; Faculty of Biology, Albert-Ludwigs-University of Freiburg, Germany; Department of Rheumatology and Clinical Immunology, Hannover Medical School, Hannover, Germany; Cellular Signalling and Cytoskeletal Function Laboratory, The Francis Crick Institute, London, United Kingdom; Immunology, Asthma and Allergy Research Institute, Tehran University of Medical Sciences, Tehran, Iran; Paediatrics Center of Excellence, Children’s Medical Center, Tehran University of Medical; Department of Genetic Medicine, Johns Hopkins University School of Medicine, Baltimore, USA Sciences, Tehran, Iran; Clinic of Rheumatology and Clinical Immunology, Center for Chronic Immunodeficiency (CCI), Medical Center, Faculty of Medicine, Albert-Ludwigs-University of Freiburg, Germany; DZIF – German Center for Infection Research, Satellite Center Freiburg, Germany; CIBSS – Centre for Integrative Biological Signalling Studies, Albert-Ludwigs-University, Freiburg, Germany; RESIST – Cluster of Excellence 2155, Hannover Medical School, Satellite Center Freiburg, Germany; Department of Infectious Disease, Imperial College, London W2 1PG, UK; RESIST – Cluster of Excellence 2155, Hannover Medical School, Hannover, Germany

## Abstract

The seven subunit Arp2/3 complex drives the formation of branched actin networks that are essential for many cellular processes including cell migration. In humans, the ARPC5 subunit of the Arp2/3 complex is encoded by two paralogous genes (*ARPC5* and *ARPC5L*), resulting in proteins with 67% identity. Through whole-exome sequencing, we identified a biallelic ARPC5 frameshift variant in a female child who presented with recurrent infections, multiple congenital anomalies, diarrhea, and thrombocytopenia, and suffered early demise from sepsis. Her consanguineous parents also had a previous child who died with similar clinical features. Using CRISPR/Cas9-mediated approaches, we demonstrate that loss of ARPC5 affects actin cytoskeleton organization and function, as well as chemokine-dependent cell migration *in vitro*. Homozygous *Arpc5*-/- mice do not survive past embryonic day 9 due to severe developmental defects, including loss of the second pharyngeal arch which contributes to craniofacial and heart development. Our results indicate that ARPC5 is important for both prenatal development and postnatal immune signaling, in a non-redundant manner with ARPC5L. Moreover, our observations add the *ARPC5* locus to the list of genes that should be considered when patients present with syndromic early-onset immunodeficiency, particularly if recessive inheritance is suspected.

## Introduction

Conserved from yeast to man (1), the Arp2/3 complex is essential for a wide variety of fundamental cellular processes through its ability to generate branched actin filament networks. Though comprised of two actin-related proteins (Arp2 and Arp3) and five additional subunits (ARPC1-5), mammalian Arp3, ARPC1 and ARPC5 are each encoded by pairs of genes (*ACTR3*/*ACTR3B, ARPC1A/ARPC1B* and *ARPC5/ARPC5L*) that give rise to proteins with 91, 67, and 67% sequence identity, respectively (2-4). Previous studies have shown that the ARPC1 and ARPC5 paralogous protein pairs confer different actin-nucleating efficiencies to the Arp2/3 complex (5), Arp3- and Arp3B-containing complexes also show divergent properties (6). This suggests that the mammalian Arp2/3 complex consists of eight iso-complexes that have evolved to perform distinct cellular or physiological functions.

In addition to the 8 different possible iso-complexes, the Arp2/3 complex itself is subject to many layers of regulation. Amongst the most important are the Wiskott-Aldrich Syndrome protein (WASp) family of Arp2/3 activators, including N-WASp, Wave1, Wave2, and Wash (7, 8). Mutations in WASP are known to cause Wiskott-Aldrich Syndrome (WAS), a complex systemic disorder combining immunodeficiency, autoimmunity, autoinflammation, atopies, and predisposition to malignancy (9-11). Since the discovery of this seminal genotype-phenotype relationship, mutations in other proteins with important roles in actin cytoskeletal regulation have been found to lead to similar human diseases; together, these are often known as “immuno-actinopathies” (12, 13).

One of the most recently identified immuno-actinopathies is the autosomal recessive ARPC1B deficiency (15, 16). Mutations in this subunit of the Arp2/3 complex result in severe inflammation and immunodeficiency, as well as impaired cytotoxic T lymphocyte maintenance and cytolytic activity [OMIM: 617718] (14-21). ARPC1A, which is normally expressed at low levels in hematopoietic lineages, is substantially upregulated in the absence of ARPC1B but unable to compensate for the latter’s loss-of-function (LOF) (15, 16, 19).

An *in vitro* system that recapitulates neonatal skeletal muscle development has also elucidated unique roles for ARPC5 isoforms (22). In this system, ARPC5L but not ARPC5 containing Arp2/3 complexes are required for the correct positioning of nuclei in the periphery of myofibers, while ARPC5 but not ARPC5L is involved in the organization of transverse triads (22). While the ARPC5 isoforms have differential functions in *in vitro* assembled myofibers, their *in vivo* physiological roles remain to be established.

We now report the consequences of ARPC5 LOF in humans and mice. We show that human ARPC5 deficiency cannot be compensated for by the presence of ARPC5L, and leads to a syndrome featuring immune disease, multiple congenital anomalies, and early postnatal death. Moreover, unlike ARPC1B mice which do not show impaired prenatal development or survival (16), constitutive ARPC5 LOF results in embryonic lethality. Thus, this gene should be included in genetic testing for families with recurrent fetal and/or infantile deaths, particularly if consanguinity is present.

## Results

### Clinical presentation

We studied the now-deceased son (V1.4) and daughter (V1.5) of consanguineous Iranian parents, who presented with multiple invasive infections and oral abscesses in early infancy (Figure 1A, Table 1 and 2). They also had two other healthy siblings – a 15-year-old sister (VI.3) and a 3-year-old brother (VI.6). Of note, the mother’s second pregnancy resulted in a first trimester miscarriage (at eight weeks); prenatal ultrasound could not identify heart formation in the fetus. V1.4 was found postnatally to have multiple congenital anomalies including a congenital heart defect (CHD) (patent foramen ovale), cleft palate, and hypoplastic corpus callosum (Table 1). He subsequently developed hydrocephalus, severe pulmonary hypertension, oral ulcers, and erosive gastritis. V1.5 was also born with CHD (moderate pulmonary stenosis and atrial septal defect) and developed oral ulcers as well as persistent diarrhea. Both patients also developed symptomatic thrombocytopenia at age 1 month (Table 2) and died of sepsis at age 3 months (Table 1). As far as we know, neither showed evidence of atopy or developed malignancy, but the increased risk for these potential complications may have been masked by their early deaths.

**Figure 1.**
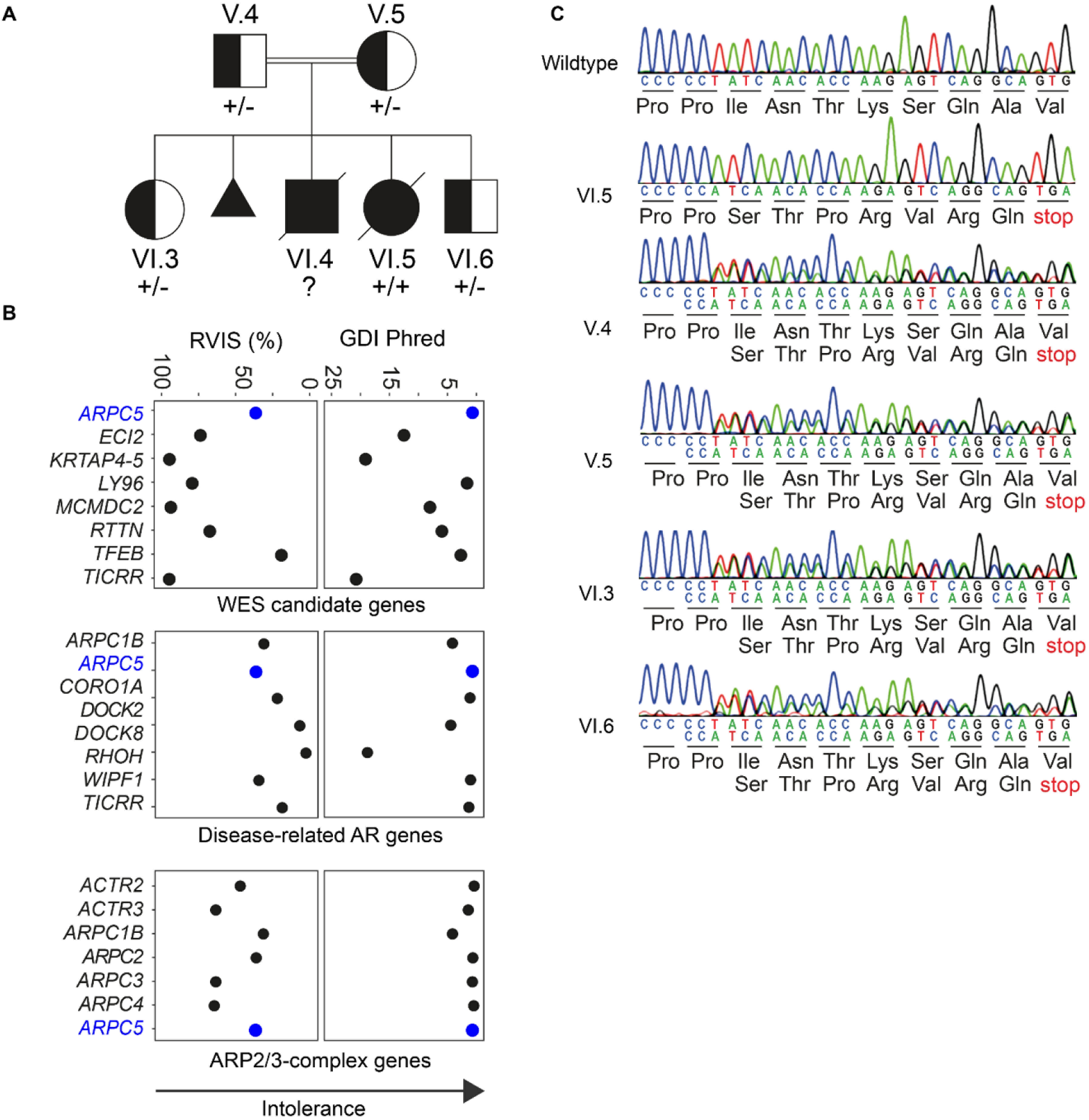
Arpc5 deficiency manifests as a severe systemic autosomal recessive disorder. (**A**) Index patients (V1.4 and V1.5) with parents and siblings. Complete family pedigree is available in Supplemental Figure 1B. Half-filled symbols represent unaffected *ARPC5* heterozygous mutation carriers. Filled symbols represent affected homozygous (and presumed homozygous) carriers. (**B**) Intolerance scores (GDI, gene damage index; RVIS, residual variation intolerance score) for genes in which homozygous variants were identified during WES analysis of VI.5 (WES candidate genes) compared to the scores for genes that are already known to be associated with IEIs (Disease-related AR genes) and related to ARPC5, or encoding other subunits of the Arp2/3 complex (ARP2/3 complex genes). (**C**) Sanger sequencing electropherograms of the index family showing the single nucleotide deletion.

**Table1:** Clinical manifestations of V1.4 and V1.5 and comparison with ARPC1B deficient patients and index patients. Estimated incidence of ARPC1B deficiency-associated features was calculated using data downloaded from GenIA (www.geniadb.net), a dedicated Inborn Errors of Immunity database that currently stores the complete literature related to ARPC1B-deficiency.

|  | Manifestation | ARPC1B-def | VI.4 | VI.5 |
| --- | --- | --- | --- | --- |
| Developmental defects | Congenital heart disease | - | Yes | Yes |
|  | Hydrocephalus | - | Yes | No |
|  | Cleft palate | - | Yes | No |
|  | hypoplastic corpus callosum | - | Yes | No |
| GI symptoms | Diarrheas | 11/30 (36.6%) | Yes | Yes |
|  | Enteritis | 4/30 (13.3%) | No | Yes |
|  | Gastritis | 2/30 (6.7%) | Yes | No |
|  | Hepatosplenomegaly | 4/30 (13.3%) | Yes | Yes |
|  | Oral ulcers | - | Yes | Yes |
|  | Gut ulcers | 2/30 (6.6%) | Yes | Yes |
| Imm. Symptoms | Recurrent infections | 26/30 (86%) | Yes | Yes |
|  | Thrombocytopenia | 14/30 (46.7%) | Yes | Yes |
|  | Sepsis | 2/30 (6.7%) | Yes | Yes |
|  | Increased IgG | 7/30 (23.3%) | - | Yes |
|  | Increased B cells | 11/30 (36.7%) | Yes | No |
|  | Reduced T Cells | 10/30 (33.3%) | Yes | Yes |

**Table 2:** Lab values of V1.4 and V1.5

| Parameter | VI.4 | VI.5 | Ref.Range |
| --- | --- | --- | --- |
| Leukocytes (cells/μl) | 6000 | 14480 | 5000-19500 |
| Mononuclear cells (%) | n.p. | 41 | 40-60 |
| Lymphocytes (%) | n.p. | 27 | 20-40 |
| Monocytes (%) | n.p. | 13 | 2-8 |
| Platelets (cells/μl * 10<sup>3</sup>) | 18 | 127 | 250-450 |
| Hb (g/dl) | 11.5 | 12.6 | 11.4-14.1 |
| IgG (mg/dl) | n.p. | 1192 | 430±119 |
| IgA (mg/dl) | n.p. | 10 | 30±11 |
| IgM (mg/dl) | n.p. | 21 | 21±13 |
| CD3 (%) | 43 | 48 | 51-77 |
| CD4 (%) | 37 | 36 | 35-56 |
| CD8 (%) | 6 | 16 | 12-25 |
| CD19 (%) | 50 | 32 | 12-36 |
| CD16 (%) | 7 | 14 | 4-18 |
| CD56 (%) | n.p. | 4 | - |

Laboratory studies for both affected siblings showed normal leukocyte counts, and reduced T-cell and platelet counts in comparison to age-matched controls (Table 2). V1.5 also had monocytosis, which may have been secondary to increased monocyte recruitment during infection; V1.4 showed increased B-cell numbers. The former also showed elevated IgG levels, while IgA was slightly decreased compared to age-matched controls (Table 2). Unfortunately, more detailed additional immunophenotyping to investigate parameters such as lymphocyte subsets, functional responses to mitogens and antigens, serum cytokine levels, or markers of type I and II interferon and/or NF-kappaB activation, was not possible due to the early deaths of both children.

### Identification of a homozygous truncating mutation in *ARPC5*

The known consanguinity of both unaffected parents was strongly suggestive of a recessively inherited defect. To confirm our hypothesis, we performed whole exome sequencing (WES) using V1.5 as proband, because V1.4 had unfortunately already passed away with no stored DNA sample available. We first interrogated genes already known to be associated with inborn errors of blood and immunity for homozygous candidate variants but identified none of interest. Application of a stringent filtering strategy (Supplemental Figure 1A) identified eight rare homozygous candidate variants (Supplemental Figure 1B). We then further narrowed down this list *via* segregation testing on available family members, performing WES on the patient’s parents, living siblings, and 14 extended family members (Supplemental Figure 2). Trio analysis found that only four of the eight candidates were heterozygous in both parents and absent in homozygosity in unaffected individuals (Supplemental Figure 1B). Amongst these, we predicted that the identified LOF variant in *ARPC5* (ENST00000359856.11:c.189delT, p.Ile64Serfs*8) could lead to a severe clinical phenotype, via potentially damaging effects on ARPC5 protein, a subunit of the Arp2/3 complex. This was supported by the recent finding that LOF in ARPC1B, another member of the Arp2/3 complex (15, 16, 21), led to some overlapping clinical features with our patients (Table 1). This variant was also notably absent from large population databases such as gnomAD exomes or genomes, TopMED/BRAVO, All of Us, Exome Sequencing Project, or the 1000 Genomes Project, and large variant databases such as ClinVar or LOVD. Moreover, it was the only gene harboring a variant with predicted high impact on protein function (frameshift). In particular, gene mutation intolerance scores based on different algorithms (GDI – gene damage index (23), and RVIS – residual variation intolerance score (24)) both found *ARPC5* to be amongst the top 1 or 2 most intolerant to functional variation of the candidate genes (Figure 1B). Importantly, unaffected family members, including the parents and unaffected elder sister (VI.3), were at most heterozygous carriers (Figure 1C; Supplemental Figure 2), while four additional relatives were also unaffected. At the time of our analysis, the mother was again pregnant and prenatal screening was performed on the fetus (VI.6), who was found to be a heterozygous carrier and carried to term (Figure 1C; Supplemental Figure 2). This son is now 3 years old and healthy, which further supports our hypothesis that biallelic *ARPC5* LOF led to the clinical problems experienced by the deceased siblings.

### *ARPC5*: p.Ile64Thrfs*8 leads to loss of ARPC5 and upregulation of ARPC5L

The homozygous variant in *ARPC5* is a single nucleotide deletion ENST00000359856.11:c.189delT, p.Ile64Serfs*8 in exon 2 predicted to result in the formation of a premature stop codon (Figure 2A). The resulting RNA transcript would be expected to undergo nonsense-mediated decay, or potentially result in expression of a truncated protein containing only the first 66 amino acids of the native 151 residue protein (ENSP00000352918.6) as well as 7 additional residues (TTPRVRQ) before the stop codon (Figure 2A). Such a truncated protein would not assemble into the Arp2/3 complex as the C-terminal half of ARPC5 is required to bind the rest of the complex (25). To examine the functional consequences of the identified variant, we targeted the same exon of the *ARPC5* gene affected by the identified c.189delT mutation via CRISPR/Cas9 in THP1 cells. We generated a c.191delT variant which also leads to a frameshift variant (p.Ile64Serfs*8) as observed in the patient with the same additional residues with the exception of the first threonine residue (STPRVRQ) (Figure 2A). This variant resulted in the loss of *ARPC5* expression both at mRNA and protein levels (Figure 2B-C). In contrast to the WT allele, transduction with lentiviral vectors expressing the ARPC5-c.189delT allele failed to rescue ARPC5 expression in ARPC5-CRISPR/Cas9-depleted (c.191delT) THP1 cells. Interestingly, upregulation of the ARPC5L isoform was seen in the absence of ARPC5 (Figure 2C). Our observations confirm that the c.189delT mutation abrogates ARPC5 mRNA and protein expression and that loss of ARPC5 leads to ARPC5L upregulation.

**Figure 2.**
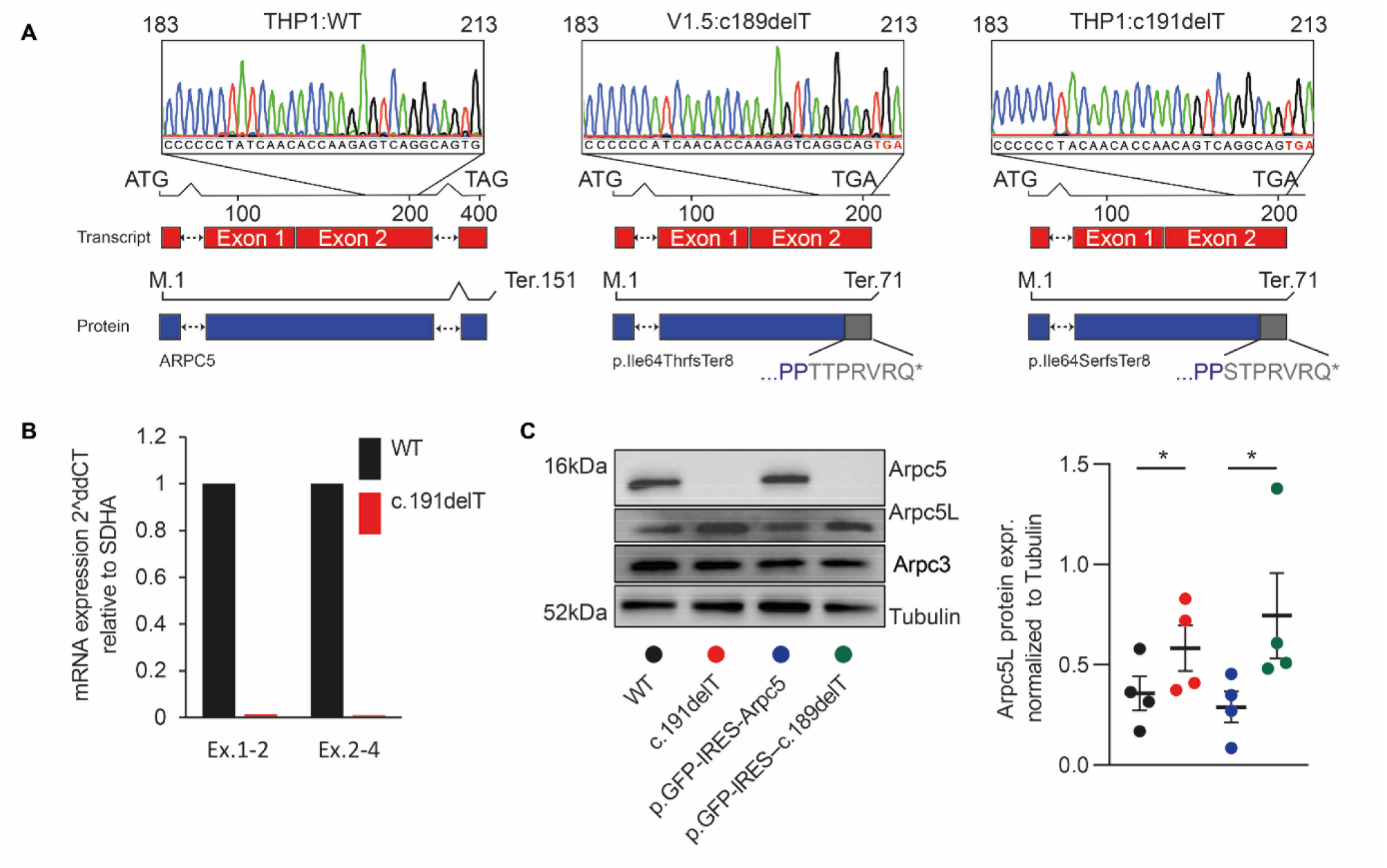
ARPC5:p.Ile64Thrfs*8 leads to loss of ARPC5 and upregulation of ARPC5L. (**A**) Schematic representation of mRNA and Sanger sequencing of gDNA respectively from WT THP1 cells, the index patient VI.5, and c.191delT THP1 cells. (**B**) mRNA quantification of ARPC5 exons 1-2 mRNA (Ex.1-2) and exons 2-4 (Ex.2-4) in WT and c.191delT THP1 cells. (**C**) Immunoblot analysis of ARPC5 and ARPC5L in WT, c.191delT THP1 cells and c.191delT THP1 cells reconstituted with WT or c.189delT ARPC5 cDNA. The graph shows the relative quantification of four independent replicates (* indicates p<0.05 as measured by a Ratio Paired T test).

### ARPC5 deficiency affects actin cytoskeleton organization and function

To investigate whether ARPC5L compensates for the lack of ARPC5, we examined the impact of ARPC5 deficiency on the organization and function of the actin cytoskeleton in HeLa cells. As observed in THP1 cells (Figure 2C), loss of ARPC5 in HeLa cells resulted in increased ARPC5L expression (Figure 3A). Nevertheless, there was a dramatic reduction in cell spreading together with a loss of actin stress fibers and focal adhesions (Figure 3B). Moreover, ARPC5-/- HeLa cells showed reduced cell migration compared to WT controls, with mean velocities and displacements of 0.17 ± 0.06 μm/min and 10.39 ± 0.94 μm vs 0.20 ± 0.06 μm/min and 15.27 ± 3.26 μm, respectively (Figure 4A). Consistent with this, ARPC5-/- cells were also significantly impaired in their ability to migrate into mechanically generated scratches relative to WT cells (Figure 4B). By 72 hours, the WT cells had achieved 100% gap closure, while the ARPC5-/- cells failed to attain even 75%. Loss of ARPC5 also delayed migration of adherent MDA-MB-231cells (a triple-negative breast cancer line) (Supplemental Figure 3A-B).

**Figure 3.**
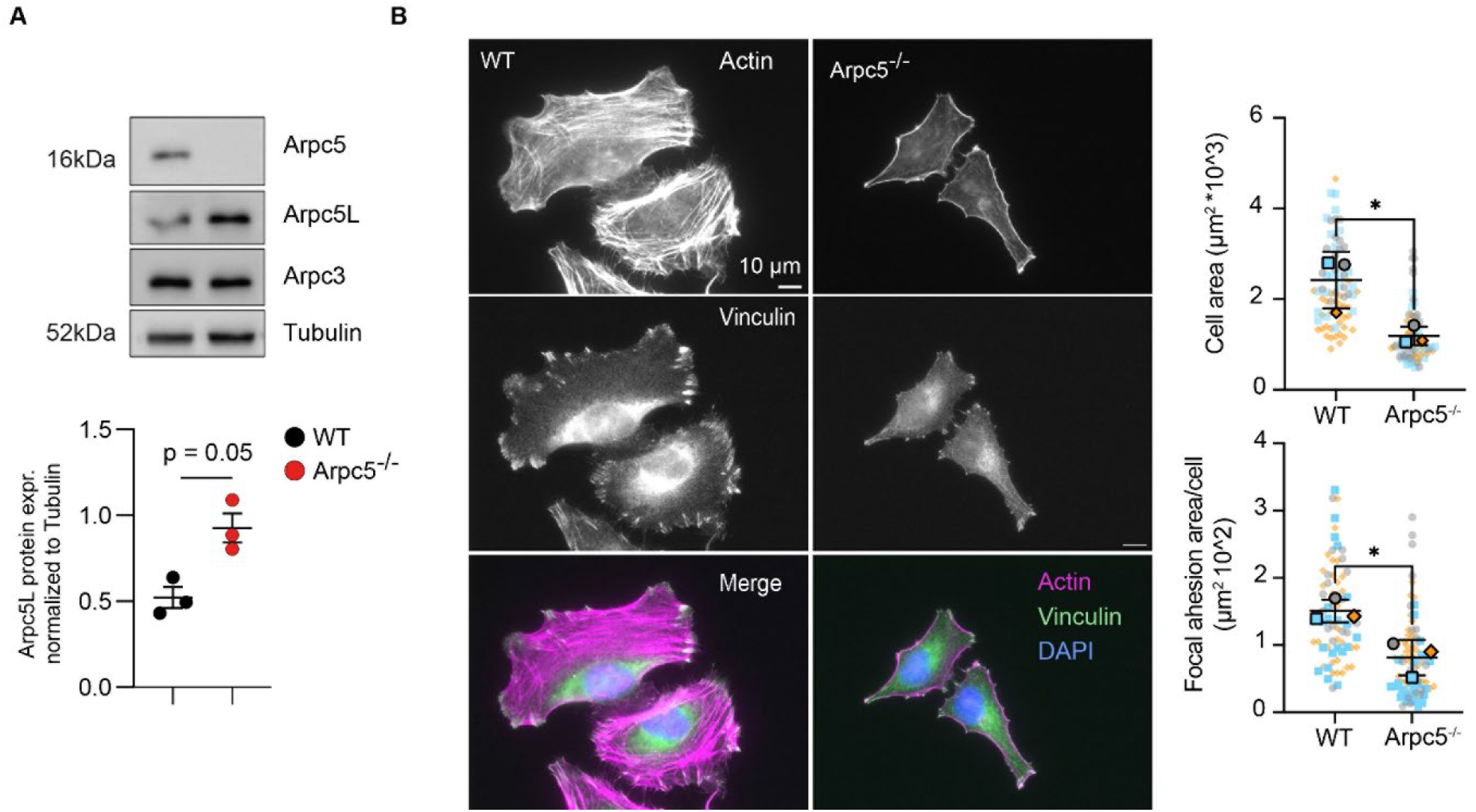
Arpc5 deficiency affects cell spreading, actin stress fibers morphology and focal adhesions. (**A**) Immunoblot analysis of ARPC5L in WT and ARPC5-/- HeLa cells, quantification and statistical analysis of independent triplicates (p=0.05 as measured by a Ratio Paired T test). (**B**) Immunofluorescence images of HeLa WT and ARPC5-/- cells labelled with Alexafluor-488 Phalloidin (Actin - Magenta), Vinculin (focal adhesions – green) and DAPI (DNA -blue). Quantification of cell and focal adhesion area of WT and ARPC5-KO HeLa cells. Three independent experiments were analyzed. n = 20-30 per experiment. Statistical analysis was performed using unpaired t-test. * indicates p < 0.05.

**Figure 4.**
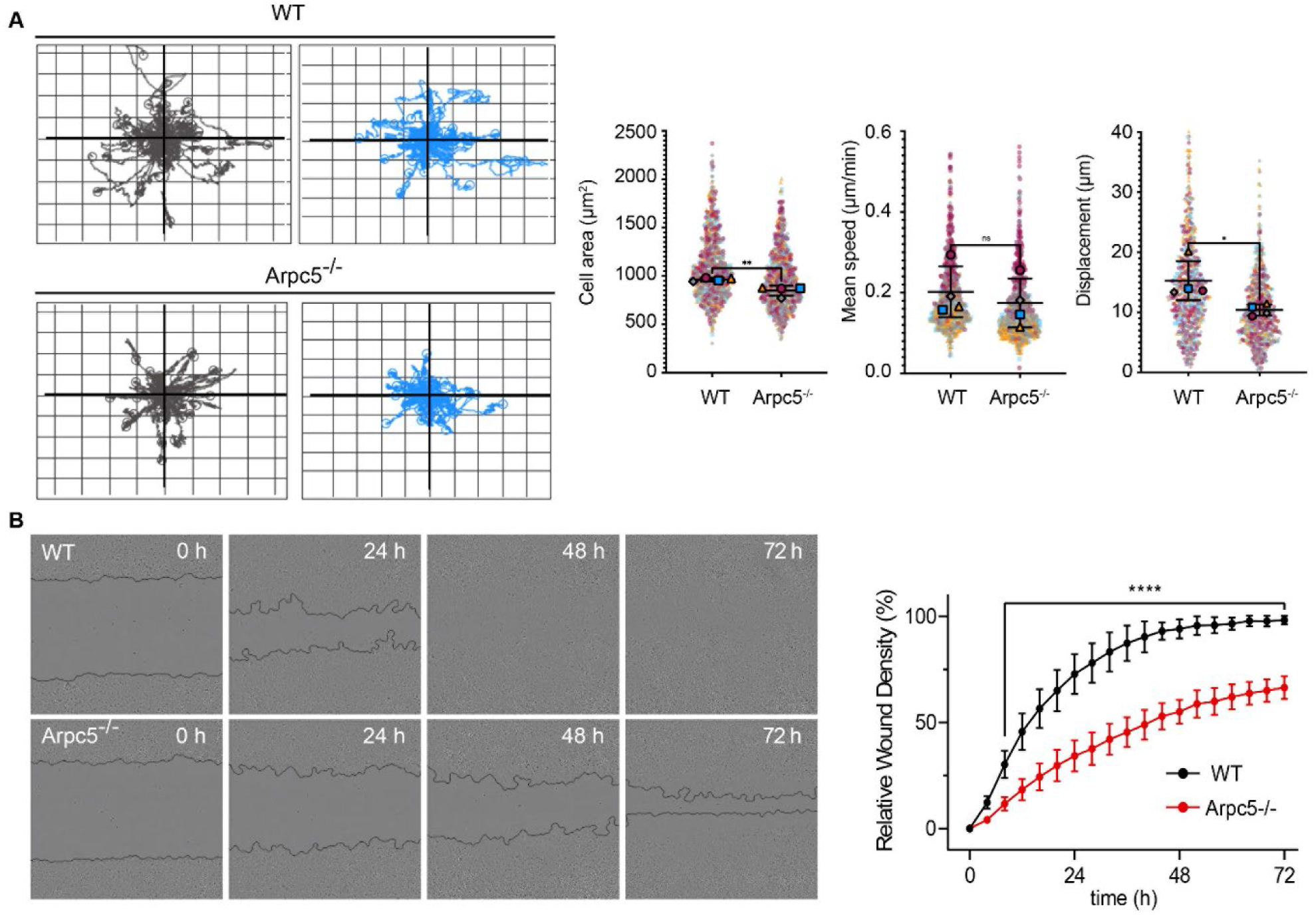
Arpc5 deficiency affects cells migration. (**A**)Two representative spider graphs illustrating the displacement of WT or Arpc5-/- Hela cells during 24-hour random cell migration. Fifty cell tracks are shown in each graph. (Right) Quantification of cell area, speed and displacement of WT and Arpc5-/- Hela cells during random migration. Four independent experiments were analysed. n > 200 per experiment. Statistical analysis was performed using unpaired t-test. * indicates p < 0.05. ** indicates p < 0.01 and ns = not significant. (**B**) Representative phase images of HeLa WT and ARPC5-/- cells are shown for the indicated times in hours after the scratch. Relative wound density of four technical replicates for each time point are shown. The statistical analysis was performed using two-way ANOVA **** indicates p < 0.0001.

Having observed defective cell migration of adherent HeLa ARPC5-/- cells, we also decided to examine the impact of ARPC5 deficiency on non-adherent Jurkat cell migration under agarose. ARPC5 deficiency was again associated with increased ARPC5L protein levels (Figure 5A). Unlike WT cells, the migration velocity of ARPC5-/- cells did not increase after stimulation with CXCL12 (Figure 5B). Interestingly there was a significant impairment in the extent of actin assembly after CXCL12 stimulation in ARPC5-/- cells relative to WT controls, though the temporal kinetics appeared to be unaffected (still peaked at 30s) (Figure 5C). These findings were also seen for the c.191delT-THP1cells (Supplemental Figure 3C). Taken together, we found that ARPC5 deficiency leads to reduced cell migration due to defects in actin organization and assembly that are not compensated for by an increase in ARPC5L expression.

**Figure 5.**
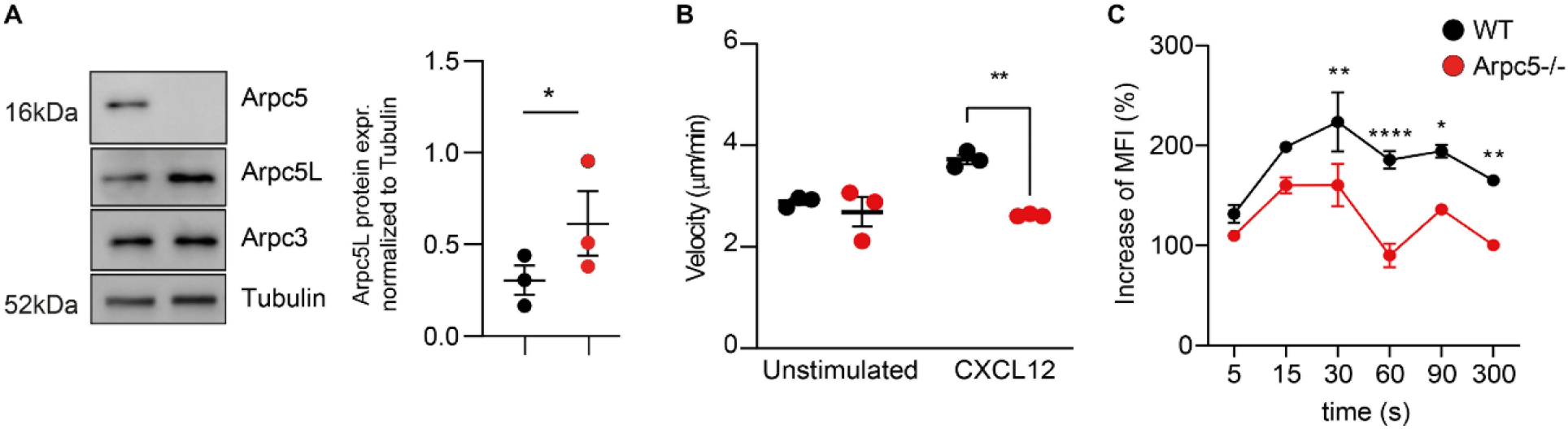
Arpc5 deficiency impairs cell migration and actin polymerization of Jurkat cells. (**A**) Immunoblot analysis of ARPC5L in WT and ARPC5-/- Jurkat cells. (**B**) Velocity of WT and ARPC5-/- Jurkat cells with and without CXCL12 treatment. Three independent experiments were analyzed. Statistical analysis was performed using Tukey’s multiple comparison test. ** indicates p < 0.01. (**C**) F-actin polymerization in Jurkat cells with or without ARPC5 at the indicated time point after stimulation with CXCL12. MFI = Mean Fluorescence Intensity and three independent experiments were analyzed. Statistical analysis was performed using two-way ANOVA. *, **, **** respectively indicate p < 0.05, p < 0.01 and p < 0.0001; error bars indicate standard deviations.

### Defective organogenesis and embryonic lethality in *Arpc5* KO mice

To further investigate and characterize the effects of ARPC5 deficiency during mammalian development, we generated *Arpc5* KO mice by targeting endogenous exon 2 with a floxed allele to inactivate the *Arpc5* gene using an early and uniformly expressed *PGK1-*Cre recombinase (Figure 6A). Based on our observations in cells, deletion of exon 2 is expected to result in loss of *Arpc5* rather than expression of a truncated protein. The genomic deletion was verified by PCR (Supplemental Figure 4). Both *Arpc5*^+/+^ and *Arpc5*^+/-^ mice developed normally through all stages until adulthood, with the correct expected frequencies. However, *Arpc5* homozygous LOF resulted in embryonic lethality as no *Arpc5*^*-/-*^ mice were identified from mating heterozygous *Arpc5*^+/-^ parents (Figure 6B). Microdissection followed by genotyping indicated that *Arpc5*^*-/-*^ embryos only become distinct from controls at 8.5-9.5 days post-fertilisation (dpf), and not earlier, and remain viable up to 9.5 dpf (Figure 6B) (26). The majority of *Arpc5* KO embryos at 8.5-9.5 dpf are seen as tissue debris, or occasionally with abnormal morphology (4 out of 26 littermates) suggesting failure of post-implantation embryonic development to progress in the absence of *Arpc5* (Figure 6C). Immunofluorescence analysis revealed that control embryos at 9.5 dpf are fully turned and have developed 19-20 somite pairs, a circulatory system including a primitive heart and the presence of blood cells, the first and second pharyngeal arches (PA1 and PA2), and the closed cranial (anterior) neuropore, which are all hallmarks of Theiler stage 14 (Figure 6D). *Arpc5* KO embryos show only 14-15 somite pairs, incomplete cranial neuropore closure, heart and PA2 formation, and have not yet completed embryonic turning. Thus, while axial patterning appears to be grossly intact, early global *Arpc5* LOF significantly disrupts tissue morphogenesis in mouse embryos, leading to post-implantation lethality.

**Figure 6.**
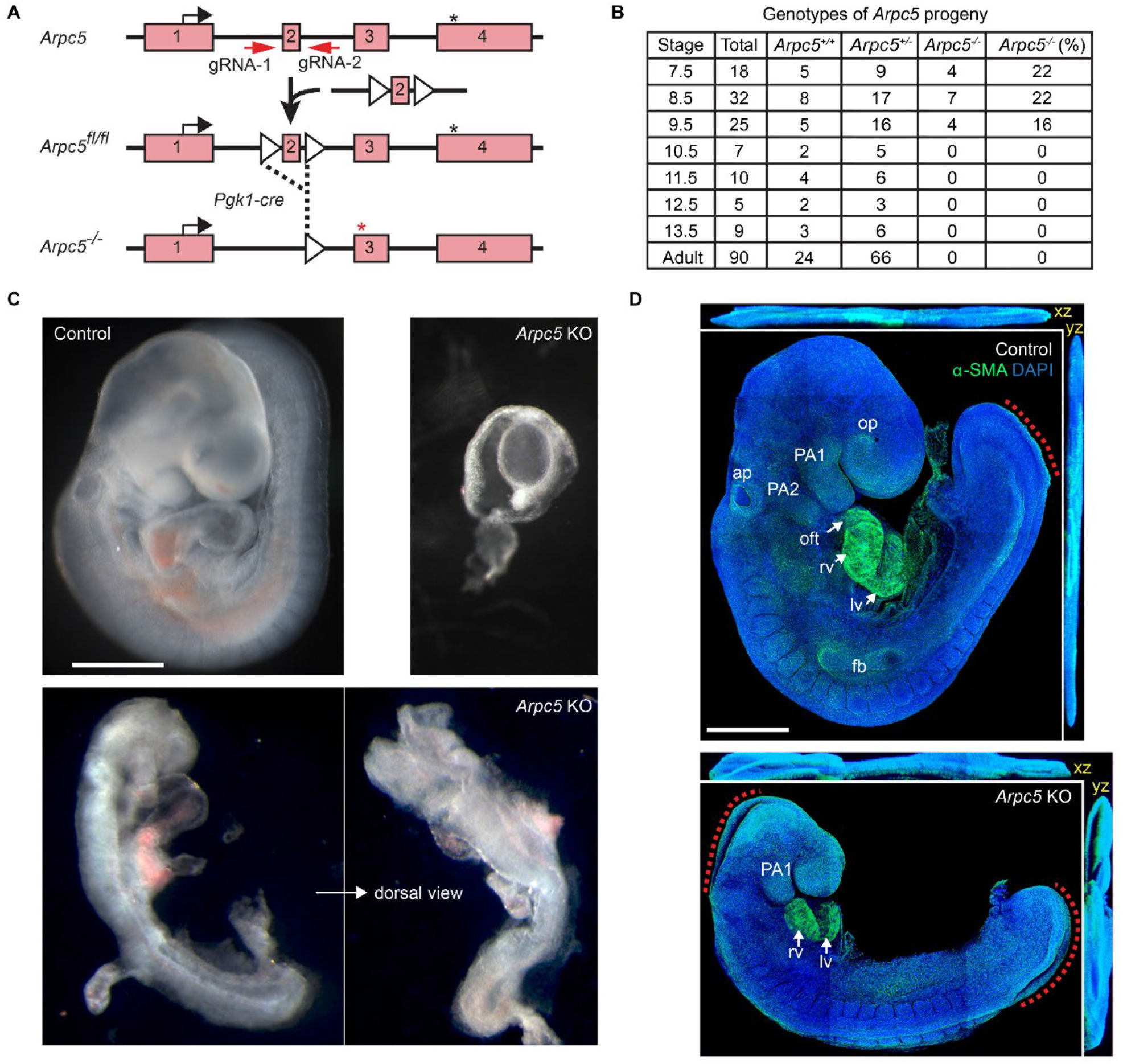
Arpc5 deficiency in mice results in defective organogenesis and embryonic lethality. (**A**) Schematic of the genomic organization of the Arpc5 locus highlighting the 4 exons (pink rectangles) as well the position of the start (black arrow) and stop (black star) codons. The strategy used to generate the conditional KO mice and the position of two loxP sites (white triangles) flanking exon 2 are indicated. Cre-mediated removal of the exon 2 results in a frameshift within exon 3 and a new stop codon (red star). (**B**) The pooled number of embryos and littermates verified as wild-type (*Arpc5*^*+/+*^), heterozygous (*Arpc5*^*+/-*^), or homozygous (*Arpc5*^*-/-*^) by genotyping from heterozygous parents. Stages indicate days post-fertilization (dpf). (**C**) Representative images of control and *Arpc5* KO embryos. Most *Arpc5* KO embryos are resorbed earlier (right) but a few show growth arrest and die by 9.5 dpf (bottom shows side and dorsal views). Scale bar, 1 mm. (**D**) Maximum intensity projection images and orthoview (xz, yz) of control and *Arpc5* KO embryos stained with alpha-Smooth muscle actin (α-SMA, green) and DAPI (blue) at 9.5 dpf. Red dotted lines indicate partial closure of spinal neuropore (control and *Arpc5* KO) and cranial neuropore (*Arpc5* KO only), respectively. The second pharyngeal arch (PA2) and the cardiac outflow tract (oft) are absent from *Arpc5* KO embryos. The first pharyngeal arch (PA1); right (rv) and left (lv) ventricular myocardium; auditory pit (ap); optic pit (op) and forelimb bud (fb) where present are indicated. Scale bar, 0.5 mm.

## Discussion

Herein, we describe human ARPC5 deficiency as a novel syndromic disorder characterized by severe, early onset infections indicative of an inborn error of immunity (IEI), along with multiple congenital anomalies involving neurological, cardiovascular, craniofacial and hematopoietic development. Although we could only perform genetic analysis on one of the two affected patients (V1.5), multiple lines of evidence support the homozygous *ARPC5* c.189delT, p.(Ile67Serfs*8) variant we identified as being causative of our index patient’s and likely also her brother (V1.4)’s severe clinical phenotypes (27): 1) The variant is sufficiently rare in control individuals, being absent from databases of either known variants or putatively health individuals, 2) It was not only found in homozygosity in a child of consanguineous parents, but also segregated appropriately with clinical presentation within a large family pedigree, 3) Multiple lines of computational evidence support its exerting a deleterious effect on the gene or gene product, 4) The variant is a predicted nullomorphic allele in a gene encoding 1 subunit of an obligate complex where LOF of another human subunit (ARPC1B) has been previously shown to lead to similar disease phenotypes and mechanisms, 5) ARPC5 deficiency affects actin cytoskeleton organization and cell migration *in vitro* and 6) ARPC5 deficiency in a mouse model is associated with developmental phenotypes involving some of the same tissues/lineages as the human disease (27).

Both genotype- and phenotype-driven inquiries identified *ARPC5* as our top candidate, but we did also consider candidate homozygous variants in 7 other genes (Supplemental Figure 1 B). While these latter were potentially consistent with our hypothetical inheritance pattern, our index patient’s presentation was less consistent with the expression and function of 6 of these genes than with that of *ARPC5*. Although human ARPC5 deficiency has never been previously reported in the literature, deficiency of ARPC1B, another subunit of the Arp2/3 complex, has been previously described – these patients show milder but similar clinical presentations as ours (15, 16, 21). In particular, both ARPC5- and ARPC1B-deficient patients share hallmarks of immunodeficiency and immune dysregulation as reflected in their histories of repeated respiratory and gastrointestinal infections, cytopenias, hypergammaglobulinemia, hepatosplenomegaly, and enteropathy.

We did not have access to patient-derived cells, but were able to use CRISPR/Cas9-based methods to model the familial variant in THP1 cells, showing that it resulted in loss of both ARPC5 mRNA and protein expression. Of note, the absence of ARPC5 impacted the organization and function of the actin cytoskeleton despite the associated up-regulation of its paralog ARPC5L. ARPC5L shares 67% homology with ARPC5 (4). However, the N-terminal half of ARPC5L is partially disordered compared to ARPC5 (25) and this structural distinction may be an important contributor to ultimate differences in Arp2/3 complex activity when assembled with one or the other of the paralogs. This paradigm has been similarly illustrated for the mammalian ARPC1 paralogs (5). Similarly to what we observed for ARPC5/ARPC5L, ARPC1A becomes substantially upregulated in the absence of ARPC1B (14-16, 21), however it is clearly insufficient to compensate for loss of the latter’s biological functions in human cells. These data and our current results provides support for distinct biological roles of these paralogs, even within the context of the same complex, confirming that the Arp2/3 complex in higher eukaryotes is actually a family of iso-complexes with different properties.

The defective embryonic turning, incomplete cranial neuropore closure, and incomplete heart and PA1-2 formation seen in *Arpc5*^*-/-*^ mice, as well as the developmental abnormalities observed in our patient, suggest a role for ARPC5 in mesoderm and neuroectoderm-derived organogenesis in higher eukaryotes. PA mesoderm cells contribute substantial parts of the pharyngeal muscles and the primitive heart (28). At 8.0 dpf the PA1 mesoderm progressively migrates into the heart tube (i.e., future ventricular myocardium), while PA2 mesoderm gives rise to the cardiac outflow tract and primitive atria during heart-looping at 8.5-9.0 dpf (Figure 6D). The migration of PA2 mesoderm towards the outflow tract are also influenced by co-migrating neural crest cells from the cranial neuropore, whose signalling may be compromised by the lack of ARPC5. Consequently, the heart and the outflow tract development in KO embryo is limited. Collectively, Arpc5 mutation in mice abrogates tissue morphogenesis such as neural tube and the heart followed by post-implantation lethality. The observed severe embryonic phenotype seen in our mouse model differs from the embryonic outcomes reported for mouse models of other core or unique subunits of the Arp2/3 complex. For example, Arp3 is required for early preimplantation embryonic development (29), and *Arpc3* LOF results in defective trophoblast outgrowth (30), while no embryonic lethality results from *Arpc1b* deficiency (16). The lack of paralogs for some members of the complex and distinct developmental expression patterns for different isoforms may both contribute to these dissimilar phenotypes, but this hypothesis awaits further experimental validation.

Moreover, there is an expanding and already significant body of evidence implicating immune dysfunction in human prenatal complications and losses. However, few definitive links have been established thus far between prenatal anomalies or susceptibility to recurrent pregnancy loss and monogenic etiologies of postnatal IEIs (31). It is not surprising that ARPC5 deficiency joins other syndromic etiologies with known multi-lineage developmental impact. However, it is interesting to note that innate or adaptive immune dysregulation resulting in autoinflammation or autoimmunity are significant and shared features across almost all the other IEIs known to be associated with significant fetal or perinatal consequences (31). Moreover, many of these other IEIs also share abnormalities of hematopoiesis with ARPC5 deficiency – most notably, gain-of-function (GOF) in SAMD9L (32), which itself has a close paralog in SAMD9. Though human disorders involving SAMD9 and SAMD9L mutations also show overlapping clinical features, they also suggest non-redundant functions for these two paralogs. Similarly, mutations in the evolutionarily and functionally related proteasomal subunits also lead to syndromic conditions featuring predominantly hematologic, inflammatory and/or neurologic phenotypes (33-36).

The observations we have presented in this paper highlight some of the most exciting emerging topics linking basic science and clinical care today. Our findings further strengthen the role of actin cytoskeletal regulation in regulation of immune responses. We further add to the emerging repertoire of monogenic causes associated with both increased risk for fetal loss and significant postnatal hematopoietic and immune problems. From a basic science perspective, our observations further our paradigmatic understanding of how paralogous human genes evolve into overlapping but distinct essential roles, via structural divergence and spatio-temporal partitioning, enabling endless functional diversity to be achieved whilst preserving an elegant molecular parsimony.

## Methods

### DNA isolation and sequencing

DNA extraction from peripheral blood samples was performed as previously described (34). Whole-exome sequencing (WES) libraries were prepared using the SureSelect Human All Exon V6 kit from Agilent (Agilent Technologies, Santa Clara (CA), U.S.A.) and sequenced in a NovaSeq 6000 system from Illumina (Illumina Inc., San Diego (CA), U.S.A). Bioinformatic analysis was performed as previously described (37). The initial strategy for variant filtering and prioritization of WES data was carried out as previously described (37), limiting the analysis to IEI-associated genes. A second (more stringent) filtering strategy was performed as shown in Supplemental Figure 2A. Segregation analysis was performed using polymerase chain reaction (PCR) after confirming the WES result. Direct sequencing was performed on an ABI 3730XL genetic analyzer (Applied Biosystems).

### Cell culture

THP1 cells (ATCC #TIB-202™) and Jurkat cells (Clone E6-1, ATCC #TIB-152™) were grown in RPMI 1640 10% FBS, 2 mM L-glutamine, 1% penicillin-streptomycin (P/S). HeLa cells (ATCC #CRM-CCL-2™) and MDA-MB-231 cells (ATCC #HTB-26™) were cultured in DMEM 10% FBS, 2 mM L-glutamine, 1% P/S. The cells were maintained at a density of 0.5×10^6^ cells/ml, changing the medium every two to three days.

### CRISPR-Cas9 knockout of ARPC5

CRISPR-Cas9 technology was used to generate ARPC5 knockout Jurkat, THP1, HeLa and MDA-MB-231 cells. The GAAAGGACGAAACACCGCTGACTCTTGGTGTTGATAGGGTTTTAGAGCTAGAAATAGCA guide targeting exon 2 of ARPC5 (https://chopchop.cbu.uib.no/) was cloned into the plasmid pMAX-Crispr with Cas9-2A-eGFP using Gibson assembly. Briefly, the vector was linearized with SwaI, the gRNA was annealed at an equimolar ratio with the universal antisense oligo (GCCTTATTTTAACTTGCTATTTCTAGCTCTAAAAC) and the annealed oligos were filled in with Klenow fragment. Finally, the digested vector and the oligo were assembled using Gibson Assembly Master Mix (NEB #E2611S). After transformation into Stbl3 chemically competent *E. coli*, single colonies were grown in LB medium and plasmid DNA was extracted. Sanger sequencing was performed for confirmation. Cells of interest were electroporated using the Neon™ Transfection System: Jurkat (10μg, 1350V, 10 ms, 3 pulses), THP1 (20ug, 1250V, 50ms, 1 pulse), HeLa (10ug, 1005V, 35 ms, 2 pulses), MDA-MB-231 (20 μg, 1400 V, 10 ms, 4 pulses). The following day, GFP cells were FACS sorted at a density of 1 cell/well in a 96-well plate in RPMI/DMEM 10% FBS, 2 mM L-glutamine. The growing clones were screened for KO by Western Blot. Clones with absent ARPC5 protein expression were confirmed by Sanger sequencing after PCR amplification of the target exon.

### Lentivirus transduction of ARPC5-KO cells

The ARPC5 WT cDNA (uniprot: O1551) and the ARPC5 cDNA containing the c.189del, p.Ile64Serfs*8 mutation (MUT) (provided by Dr. Antonio Carusillo), were cloned into pLenti-IRES-GFP-Puro plasmid. HEK 293T cells were transfected with pLenti-ARPC5 WT/MUT-IRES-GFP-Puro, psPAX2, and pMD2G to produce lentiviral particles. After 48h, the supernatant was transferred to ARPC5 KO Jurkat, THP1, HeLa and MDA-MB-231 cells. Cells were incubated for 48 and sorted for GFP expression.

### SDS-PAGE and Western Blotting

Cells were collected and lysed in 2x RIPA buffer + 2x Protease Inhibitors (Roche #4693159001). The protein concentration was determined with the Pierce™ BCA Protein Assay Kit (ThermoFisher #23225) according to the manufacturer’s protocol. Equal amounts of lysates were loaded on a 14% or 18% SDS-Page for size fractionation. After electrophoresis, proteins were immunoblotted on a PVDF-membrane (Sigma-Aldrich #3010040001) for 1h at 100V and blocked with 5% milk in TBS and 1% Tween20. Proteins were detected with rabbit anti-ARPC5 (Novus #NBP2-67350), rabbit anti-ARPC5L (Abcam #ab169763) or rabbit anti-ARPC3 (anti-Arp2/3 Complex Antibody Clone 13c9 Millipore #MILL-MABT95). HRP-conjugated goat anti-rabbit IgG (Cell Signaling #7074S) was used as a secondary antibody before detection with ECL chemiluminescent substrate (Cell Signaling #12630S). Anti-Tubulin (Proteintech #HRP-66031) was used as a loading control and detected as a 55 kDa band.

### Immunofluorescence Analysis

Cells were seeded on coverslips in a 12-well plate and incubated overnight in RPMI 1640 supplemented with 10% FBS and 1% P/S. Cells plated on coverslips were incubated with 4% paraformaldehyde for 10 minutes at room temperature and washed three times with 1x PBS. Cells were then incubated with 0.5% Triton X-100 for 2 minutes at room temperature and washed three time with 1x PBS. Coverslips were incubated with blocking buffer for 30 minutes at room temperature. Coverslips were incubated with mouse anti-vinculin antibody (Sigma V4505, 1:500) overnight at 4°C. Coverslips were then washed three times with 1x PBS and incubated with Alexa Flour 488-conjugated anti-mouse antibody (ThermoFisher A21202, 1:500) and Alexa Flour 647-conjugated phalloidin (Thermo Fisher A22287, 1:500) for 30 minutes at room temperature. After three washes in 1x PBS, cells were incubated with DAPI (CST 4083S, 300 nM) for 5 minutes. Coverslips were then washed twice with 1x PBS and once with distilled water before mounting the coverslips onto microscopy slides using 4 μL Mowiol. Coverslips were imaged on a Zeiss Axioplan2 microscope with a 63 x/1.4 NA Plan-Achromat objective and a Photometrics Cool Snap HQ cooled charge-coupled device camera. Data for 20-30 cells per sample were analyzed using FIJI.

### Random migration assay

Hela WT and ARPC5-KO cells were seeded in 24-well glass bottomed plates (Cellvis) at 1*10^4^ cells/well in 500 μL RPMI 1640 supplemented with 10% FBS and 1% P/S. Cells were imaged and analyzed using the Livecyte system (Phasefocus) every 10 minutes for 24 hours at 10x with 2 fields of view per well. The imaging chamber was maintained at 37°C with 5% CO_2_.

### Wound healing assay

HeLa and MDA-MB-231 WT and ARPC5-KO cells were seeded in a 96-well ImageLock plate (Sartorius #4806) and incubated for 24 h. The following day, cells were scratched using the Incucyte® WoundMaker 96 (Sartorius). Cells were washed twice with PBS and cell migration towards the scratch was monitored with Sartorius Incucyte S3 System for 72 h, with one image obtained per hour under brightfield 10x magnification. Images were analyzed using IncuCyte 2019B Rec2 software. The relative scratch intensity was calculated as %RWD(t) = [(wt-w0)/(ct-c0)]·100

### Under agarose migration assay

Glass-bottomed 8-well μ-slides (ibidi) were coated with human ICAM-1 (3 μg/mL; R&D system) overnight at 4°C. The wells were then washed and blocked with 2% fatty acid-free BSA (Sigma-Aldrich) for 20 minutes at room temperature. Agarose gel mixture was prepared by mixing 2x phenol red-free HBSS, RPMI supplemented with 20% FBS + 2% P/S + 4 mM L-glutamine + 2x NEAA + 2x sodium pyruvate + 50 mM HEPES + 50 μM β-mercaptoethanol, and 2% liquid UltraPure Agarose (Thermo Fisher) in 1:2:1 ratio. The mixture was shaken at 300 rpm at 56°C for 10 minutes. Prior to adding the mixture to the wells, human CXCL12 (250 ng/mL; Peprotech) or vehicle control was added to the gel mixture. Gel mixtures were then added to the wells accordingly and allowed to solidify and cool for at least one hour. Jurkat WT and ARPC5-KO cells were labelled with CTV or CFSE (Thermo Fisher) according to manufacturer’s protocol prior to imaging. Propium iodide was added at 3 μM to identify dead cells. Approximately 6*10^4^ cells in 10μL were injected into an agarose gel matrix. Cells were then imaged using an inverted widefield Nikon Ti2 Eclipse long-term time lapse system with an LED illumination system. The imaging chamber was maintained at 37°C with 5% CO_2_. A Nikon 20x Ph2 planar (plan) apochromatic (apo) (0.75 N.A.) air objective was used to acquire images at a single z-slice every 15 s for 10 min with 3 fields of view per well. Cell shape and migration parameters were analyzed using FIJI plug-in TrackMate.

### CXCL12-induced F-actin polymerization assay

Jurkat and THP1 WT and ARPC5 KO cells were seeded in 96-well plate and stimulated with RPMI 1640 + 100 ng/ml CXCL12 (Peprotech, 300-28A) or RPMI 1640 for the indicated time points. Cells were fixed with ice-cold 4% PFA for 20 min and permeabilized with 0.1% Triton-X for 10 min. To detect F-actin, cells were incubated with AlexaFluor647-Phalloidin (Cell Signaling #8940S) for 20 min. Median fluorescence intensity (MFI) was measured using BD LSRFortessa™ and data was analyzed using FlowJo (Version 10.8.1). Increase in MFI was calculated as a percentage using the following formula, with MFI of the unstimulated samples defined as 100% (35): 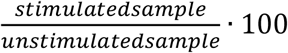

### Generation of conditional mouse models

Mice were bred and maintained in a specific pathogen-free room with a 12-hour light/dark cycle, and access to ad libitum food and water. The conditional *Arpc5* knockout (KO) strain was generated by the Crick Genetic Modification Service using a CRISPR-Cas9 approach in embryonic stem cells (ESC) with gRNA-1, 5’-TAGTTTCAGTATAAAGTCTA-3’, and gRNA-2, 5’-CCCTTTAAAGCTGGACGTGG-3’, which introduce the two 5’ and 3’ loxP sites within intron 1-2 and intron 2-3, respectively. Removal of exon 2 induces a frameshift and an exogenous stop codon within exon 3, resulting in only the first 55 residues of ARPC5 being expressed. The two Cas9-guide RNA (gRNA)-Puro plasmids were generated by inserting the CRISPR-Cas9 target sequences into PX459 plasmid (a gift from Feng Zhang (38)). #48139 Addgene, Watertown, MA, US). The pMA donor plasmid contains a synthesized 734 bp fragment corresponding to exon 2 floxed by two loxP sites and 1 kb flanking homology arms (GeneArt, Thermo Fisher Scientific, Waltham, MA, USA). Sequence-verified Cas9-gRNA-Puro plasmids and pMA donor plasmid were co-transfected into in-house C57BL6/N 6.0 ESC using Lipofectamine 2000 (Thermo Fisher Scientific, Waltham, MA, USA) prior to selection with 2 μg/ml puromycin for 48 hours. Putative targeted clones were initially identified by quantitative real-time PCR assays on genomic DNA (gDNA) using gene-specific probes designed by Transnetyx (Cordova, TN, USA) to detect the integration of 5’ and 3’ loxP sites, respectively. Integrity of the loxP integrations was confirmed by Sanger sequencing of PCR amplicons. Correctly targeted ESC clones were microinjected into blastocysts derived from albino C57BL/6J (B6(Cg)-Tyrc-2J/J; strain #000058, The Jackson Laboratories, Bar Harbour, MN, US), and then transplanted into the uteri of pseudo-pregnant CD1 females. The resulting chimeras were crossed to albino C57BL/6J, and their offspring were initially screened for the integration of the loxP sites by amplicon sequencing. Heterozygous floxed offspring were validated by Sanger sequencing across a 3.2kb amplicon encompassing the targeted allele. Two founder strains were maintained on a C57BL/6J background by backcrossing more than three times. Constitutive *Arpc5* KO mice were obtained by mating Arpc5 floxed mice with *Pgk1-cre* (B6.C-Tg(Pgk1-cre)1Lni/CrsJ (39). All mice were genotyped at weaning age at Transnetyx as described above.

### Study Approval

All animal work was authorised by UK Home Office project licences P7E080263 and personal licences following the approval by Animal Welfare and Ethical Review Body of The Francis Crick Institute. Patients were registered in the Iranian Primary Immunodeficiency Registry (IAARI IPIDR) at the Immunology, Asthma & Allergy Research Institute (IAARI). The Ethical Committee of IAARI approved this study (#IR.TUMS.IAARI.REC.1397.002), and written informed consent for participating in this study was obtained from their parents

### Whole-mount embryo immunofluorescence and volume imaging

Timed-pregnant females were used to harvest staged embryos based on the date of vaginal plug as 0.5 days post-fertilisation (0.5 dpf). Embryos up to 13.5 dpf were dissected to examine the morphology under Leica MZ16 stereomicroscope (Leica Microsystems, Wetzlar, Germany) equipped with MicroPublisher 6 colour camera (Teledyne Photometrics, Tucson, AZ, USA). The yolk sacs were harvested for genotyping by PCR with the following primer pairs: Arpc5-1 For, CCAGAATAATCAGCCAGCATTTCAG, and Arpc5-1 Rev, CTTGTCACAAGCTCCCTTTAAAGC, which generates an 819 bp unexcised product and 119 bp excised product for the Arpc5 allele as well as Cre For, TCATCTCCGGGCCTTTCG, and Cre Rev, GACAGAAGCATTTTCCAGGTATGC, which yields a 200 bp product for Cre transgene. Dissected embryos were fixed with cold 4% paraformaldehyde (PFA) in PBS overnight at 4 °C. The embryos were washed with PBS containing 0.1% Triton X-100 (PBST) three times for 1 hr each, and then incubated with PBST containing 2% BSA (blocking buffer) for 4 hr at room temperature. The embryos were subsequently incubated overnight at 4 °C in solution containing Cy3-conjugated α-smooth muscle actin monoclonal antibody (1:100, C6198; Merck, Darmstadt, Germany), Alexa647-Phalloidin (1:200, Thermo Fisher Scientific, Waltham, MA, USA), and DAPI. Embryos were washed three times with PBST for 1 hr and mounted in 1% Agarose solution before imaging. Z-stack tiled images were obtained using an inverted Zeiss 880 confocal microscope and Plan-Apochromat 10x/0.45 M27 air objective at 1024 × 1024 pixels with a step of 5 μm. The images were stitched and adjusted for intensity using Zeiss Zen Blue 2.6 software.

### Statistical Analysis

Statistical significance was calculated with, a paired ratio t-test, an unpaired t-test, Two-way ANOVA or One-way ANOVA with Tukey’s correction using GraphPad Prism 6.0 software. A p value of <0.05 was considered statistically significant (*p<0.05; **p<0.01; ***p<0.001; ****p<0.0001.

### Reverse Transcription Quantitative Polymerase Chain Reaction (RT-PCR)

Total RNA from cells was extracted using the RNeasy Mini Kit (Qiagen) according to the manufacturer’s instructions. Complementary DNA (cDNA) was generated using 1 μg RNA using M-MLV Reverse Transcriptase (Promega) according to the manufacturer’s protocol. Quantitative RT-PCR was performed using SYBR Green Dye (Applied Biosystems) on the StepOne Plus (Applied Biosystems). Results were normalized to the tissue housekeeping gene SDHA. Relative expression levels were calculated as 2^(Ct (gene) − Ct (SDHA))^.

## Author Contributions

Study design: MW, MP Experiments: ES, NK, SH

Analysis and Interpretation: ES, NK, SH, MP, AC

Writing: MP, ES, AC and NK prepared an initial draft, which XP expanded upon significantly, and then all authors contributed to revisions and generation of final text.

Figure preparation: ES, NK, SH, AC, MP

All authors reviewed and approved the manuscript.

## Acknowledgements

We thank Dr. Antonio Carussillo from the laboratory of Dr. Claudio Mussolino at the Institute for Transfusion Medicine and Gene Therapy at Center for Translational Cell Research, Germany for providing us with the ARPC5 WT and ARPC5 c.189delT cDNA. Thanks to Dr. Marie Follo from the Light Core Facility Unit for her support and teaching of the microscopes and Jan Bodinek for cell sorting and his support with the flow cytometers. We thank Dr. Ian Rosewell and Jessica Olsen in the Genetic Modification Service facility at the Crick for generating *Arpc5* floxed mice as well as the Biological Research Facility for animal maintenance. We also thank Drs Caetano Reis e Sousa and Carola Vinuesa (Crick Instititute) for providing comments on the manuscript.

## Funding Acknowledgements

MP, BG and AC were funded by the Deutsche Forschungsgemeinschaft (DFG, German Research Foundation) under Germany’s Excellence Strategy — EXC 2155 — project number 390874280. MP is also funded by the Deutsche Forschungsgemeinschaft (DFG, German Research Foundation) – TRR 359 – Project number 491676693 and by the Fritz Thyssen Foundation (grant number: 10.18.1.039MN). NK, SH and MW were supported by the Francis Crick Institute, which receives its core funding from Cancer Research UK (CC2096), the UK Medical Research Council (CC2096), and the Wellcome Trust (CC2096) as well as by additional funding from the European Research Council (ERC) under the European Union’s Horizon 2020 research and innovation programme (grant agreement No 810207 to MW).

## Figures and figure legends

**Supplemental Figure 1.**
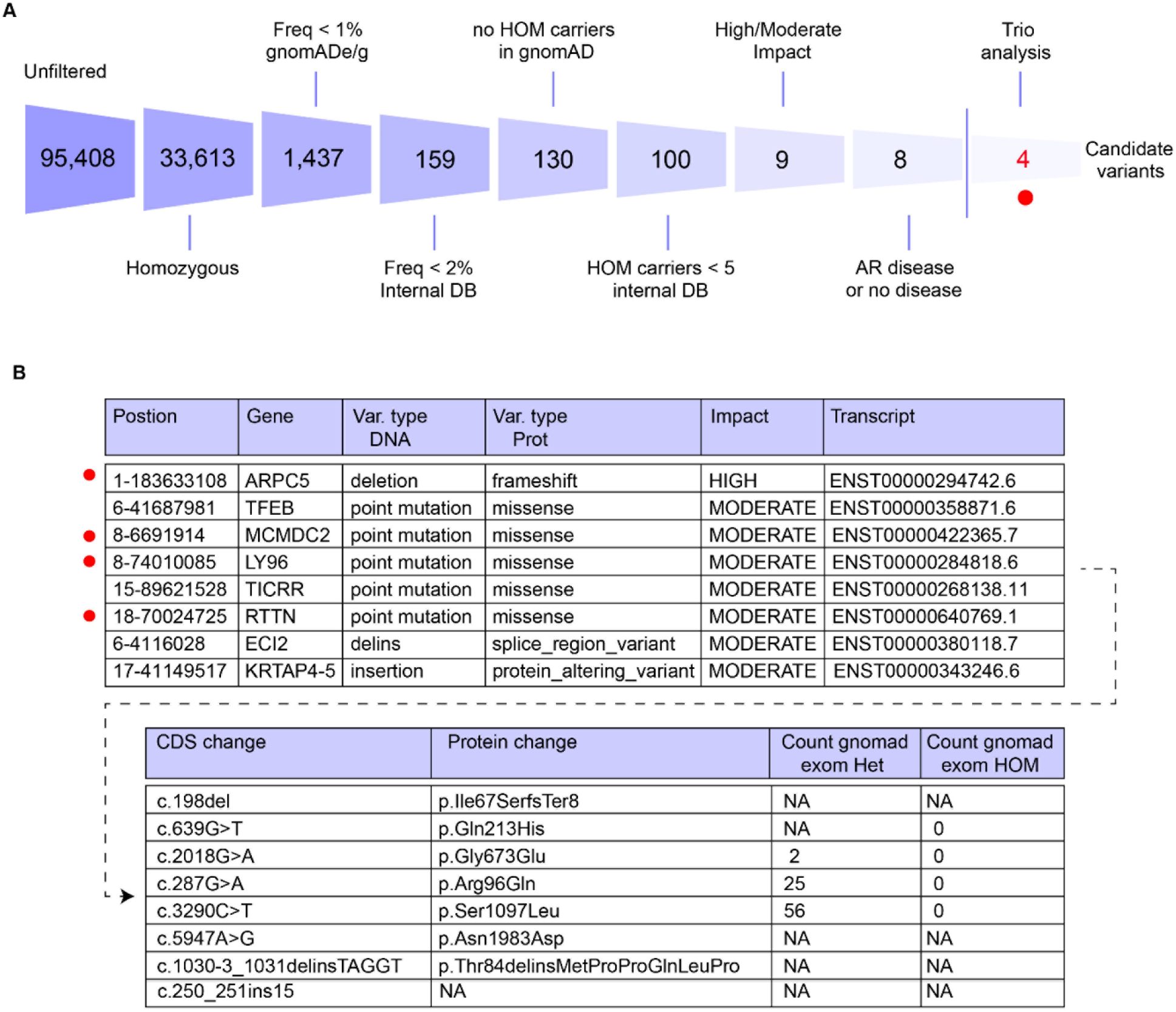
WES variant filtering strategy for index patient V1.5. (**A**) Diagram of the filtering strategy employed for the WES analysis of the index (VI.5) patient (AF = Allele Frequency; 1KG = Thousand Genome Project, https://www.internationalgenome.org/; GnomAD = Genome Aggregation Database, https://gnomad.broadinstitute.org/; (**B**) List of homozygous rare variants present in VI.4 and heterozygous in both parents. This list was filtered as detailed in the Material and Methods.

**Supplemental Figure 2.**
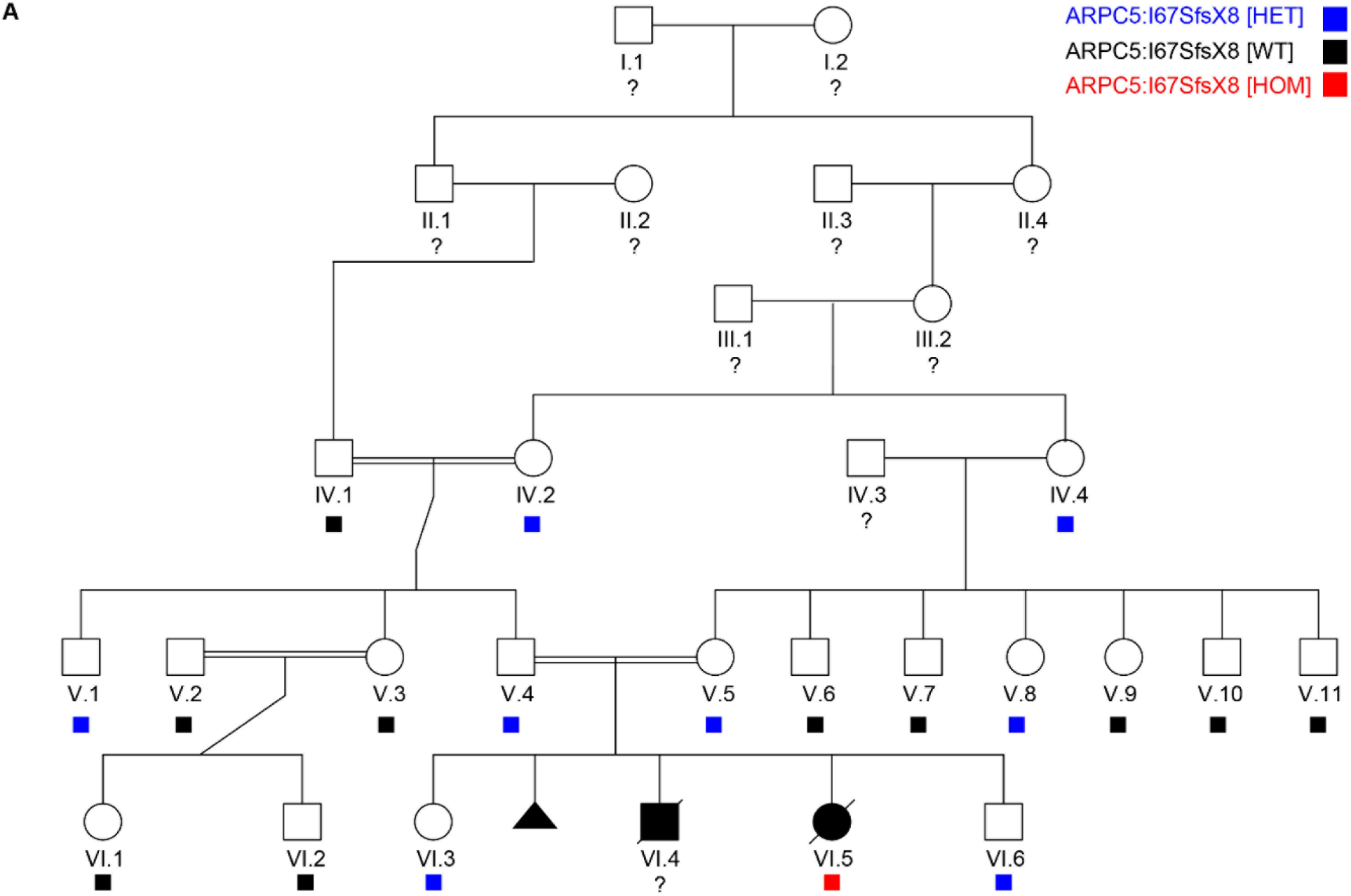
Extended index family pedigree showing *ARPC5* variant segregation. Pedigree of the extended family showing the segregation of the c.189delT variant in *ARPC5* identified in VI.5.

**Supplemental Figure 3.**
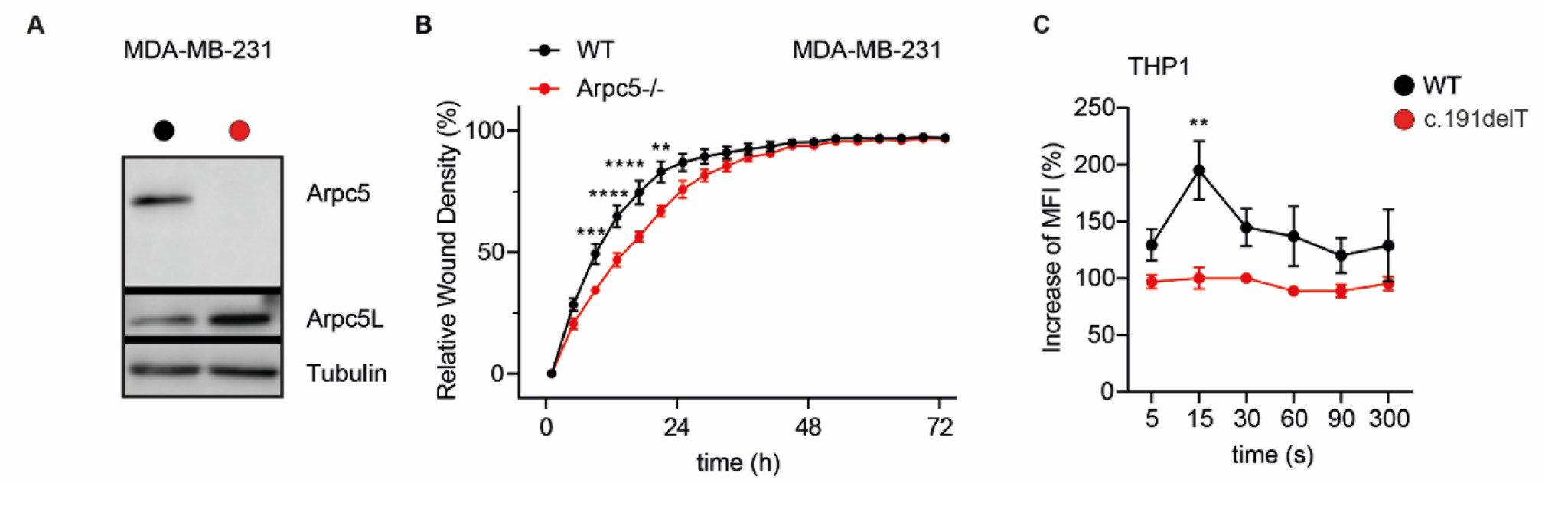
ARPC5 deficiency affects actin cytoskeleton organization and function. **(A)** Confirmation of ARPC5 deficiency in MDA-MB-231 cells via Western Blot. **(B)** Relative wound density of MDA-MB-231 WT and *ARPC5-/-* cells. (**C**) F-actin polymerization in c.191delT THP1 cells with or without ARPC5 at the indicated time point after stimulation with CXCL12. Three independent experiments were analyzed. Statistical analysis was performed using a Two-way ANOVA. **, ***, **** indicates respectively p < 0.01, p < 0.001 and p < 0.0001; error bars indicate standard deviations; MFI = Mean Fluorescence Intensity.

**Supplemental Figure 4.**
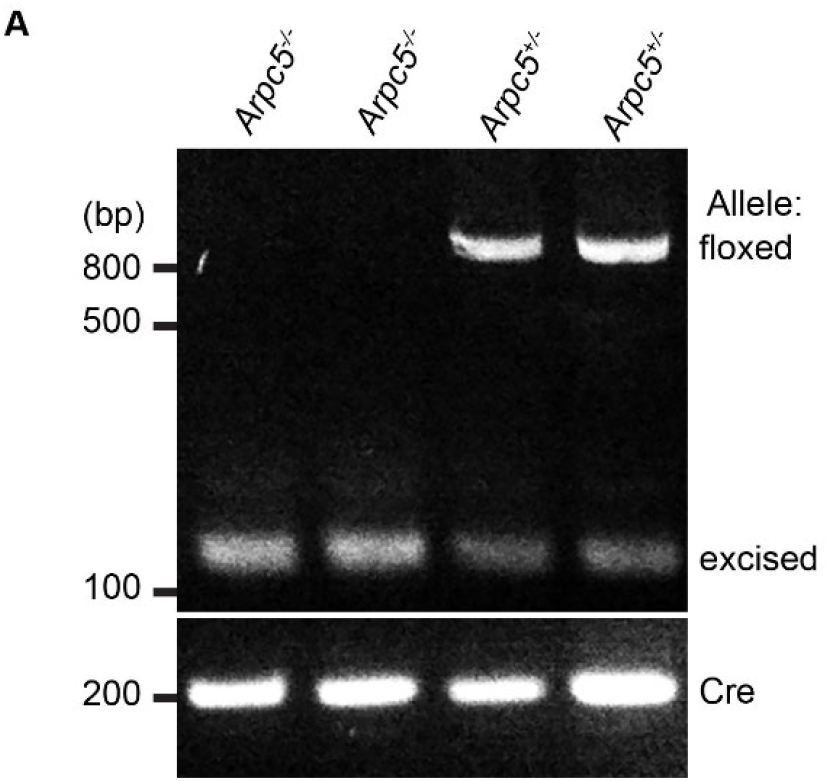
Arpc5 deficiency in mice results in defective organogenesis and embryonic lethality. Genotyping PCR showing the floxed and excised allele of *Arpc5*, as well as *Cre* transgene.

